# Estradiol Reprograms Microglia to Create an Immune-Suppressed Niche Permissive to Breast Cancer Brain Metastasis

**DOI:** 10.64898/2026.02.20.707087

**Authors:** Karen L.F. Alvarez-Eraso, María J. Contreras-Zárate, Andrew Goodspeed, James Costello, Jenny A. Jaramillo-Gómez, Stella Koliavas, R. Alejandro Marquez-Ortiz, Morgan S. Fox, D. Ryan Ormond, Peter Kabos, Mercedes Rincon, Diana M. Cittelly

## Abstract

**Background:** Young age is an independent risk factor for the development of breast cancer brain metastases (BM). Prior work showed that 17β-estradiol (E2), the predominant premenopausal hormone, promotes BM of tumors intrinsically unresponsive to E2, in part through modulating estrogen receptor-alpha expressing (ERα⁺) glial cells. However, how E2 reshapes the brain tumor microenvironment (TME), particularly microglia-mediated immunity, and its impact to BM progression remains unclear.

**Methods:** scRNA sequencing and multiparametric flow cytometry were used to define the impact of E2 and E2-suppression on brain immune-cell populations across different stages of BM progression using spontaneous and experimental models of BM. Depletion of microglia and T cell co-cultures were used to study microglia’s role in E2-induced BM. The effects of E2-suppression alone or in combination with whole brain radiotherapy were tested in preclinical models mimicking late-stage BM.

**Results:** E2 repressed immune surveillance and immune activation programs in microglia from early to late stages of brain metastatic progression, suppressing recruitment of effector immune cells to BM. Estrogen suppression, in turn reactivated anti-tumoral signaling in microglia and increased recruitment of effector immune cells to the brain. Microglia from E2-stimulated BM-bearing mice showed decreased ability to induce interferon cytotoxic function and expansion of activated T cells. Conversely, E2-suppression reactivated an effective anti-tumoral response and synergized with RT to significantly decrease BM progression.

**Conclusion:** These findings reveal a previously unrecognized mechanism by which E2 accelerates BC-BM progression through microglial immunosuppression and support evaluation of endocrine therapies as adjunct treatments for ER⁻ brain metastases.

**Importance of the Study:** Standard of care for BM includes stereotactic radiosurgery (SRS) alone or in combination with surgery, systemic chemotherapy or targeted therapies. Our studies show that ovariectomy (which eliminates ovarian E2) and aromatase inhibitors (AIs, which eliminate peripheral E2 synthesis) reduce progression of BM when used in combination with WBRT and in immuno-competent models. We demonstrate E2 promotes an immunosuppressive brain microenvironment from early stages of metastatic progression, in part through modulation of myeloid cells and repression of recruitment of effector T cells to the brain. Thus, these studies suggest that FDA-approved E2-depletion therapies (aromatase inhibitors and selective-estrogen modulators) could be used in combination with brain irradiation to decrease BM progression.

## INTRODUCTION

Breast cancer (BC) is the second most common primary tumor responsible for BM, particularly in women with HER2^+^ and triple-negative (absence of E2 receptor (ER^−^), progesterone receptor (PR^−^), and HER2 amplification) tumors^1–3^. Current treatment options for BM (including surgery, radiation, and chemotherapy or targeted therapies) have limited success and often worsen neurological function^4^. In fact, women with brain metastatic BC face a poor prognosis, with median overall survival ranging from 4 to 9 months after BM diagnosis, with outcomes varying across BC subtypes^5,6^. The high incidence of BM in triple-negative and HER2^+^ tumors has been attributed to the intrinsic aggressive nature of these subtypes. However, young age (defined as <50 years old, representing peri- or pre-menopausal woman) is an independent risk factor for the development of BM *regardless of breast tumor subtype*^7,8^, suggesting that tumor extrinsic mechanisms impact the progression of BM in younger women.

E2, the predominant pre-menopausal hormone, plays distinctive roles in brain function that can promote brain metastatic progression. E2 acts through nuclear and membrane-bound ERs that are expressed in neurons and glial cells throughout different brain compartments^9,10^, and both classical and non-classical E2 signaling pathways are intertwined in mediating neuroprotective and homeostatic processes^11,12^. In addition to ovarian E2, specific areas of the male and female brain synthesize E2 from testosterone precursors through the action of the aromatase enzyme^13–15^. We have previously shown that pre-menopausal levels of E2 promote brain metastases of tumors intrinsically unresponsive to E2, and that ovariectomy (OVX, which eliminates ovarian E2) in combination with aromatase inhibitors (AIs, which eliminate brain and peripheral E2 synthesis) *prevented brain metastatic colonization*^16,17^. Yet, the potential use of endocrine therapies in the management of BM is hindered by an incomplete understanding of how E2 alters the brain tumor-microenvironment (TME) and whether targeting the changes induced by E2 are sufficient to impact therapeutic responses.

Microglia (the brain-resident macrophages) express ERs (both ERα and ERb) and are the first responsive cells after cancer cell extravasation^18,19^, mediating the neuroprotective and homeostatic effects of E2 in the brain^20,21^. Several studies have established the important and often dual role of microglia to influence the progression of primary and secondary brain tumors^22,23^, as well as their crosstalk with T cells in determining tumor outcomes^24,25^. E2 has been shown to suppress microglia functions in certain brain pathologies and is well documented as a potent immunosuppressant that dampens macrophage inflammatory responses while promoting an alternatively activated, anti-inflammatory, tissue-repair phenotype^26^. Consistent with these findings, E2-stimulated macrophages have been shown to promote melanoma growth by adopting a pro-tumoral polarization that suppresses CD8⁺ T cell activity and enhances T cell exhaustion^27^. However, the extent to which E2 could modulate microglia function in the context of BC-BM remains unknown.

Here, we demonstrate that E2 represses immune surveillance and immune activation programs in microglia from early to late stages of brain metastatic progression, suppressing recruitment of effector immune cells to the BM. E2 suppression, in turn, reactivates anti-tumoral signaling in microglia and restores recruitment of effector immune cells to the brain. Moreover, we demonstrated that microglia from E2-stimulated BM-bearing mice have decreased ability to induce interferon cytotoxic function and expansion of activated T cells, thereby diminishing the anti-tumorigenic T cell immune response. Since BM are usually treated with brain radiotherapy (RT), either as whole brain radiotherapy (WBRT) or stereotactic radiosurgery (SRS), we show that OVX in combination with the AI letrozole increases the effectiveness of radiation in decreasing BM progression. These studies provide a novel mechanism whereby E2 promotes rapid progression of BM and provides a rationale for the clinical testing of endocrine therapies in the management of BM from ER^−^ tumors.

## RESULTS

### E2 influences microglia activity and suppresses effector immune cell recruitment in the brain pre-metastatic niche

As microglia are central regulators of neuroinflammatory responses in the brain, we sought to determine how E2 modulates microglia function and influences immune cell recruitment during BM progression of BC cells that are intrinsically unresponsive to E2 (**Suppl. Fig 1A, B**). For this, we performed multi-parametric flow analysis of brain immune cells isolated from E2-treated or E2-depleted mice at different stages of brain metastatic colonization. First, we interrogated the early stages of brain metastatic dissemination (here referred to as pre-metastatic niche) using a spontaneous model of BM. For this, E2-unresponsive EO771BR1-GFPLuc+ cells were implanted orthotopically in the MFP of OVX-female C57BL/6 mice supplemented with premenopausal levels of E2, Veh, or letrozole (**Fig 1A**). Mimicking the natural progression of BC, disseminated BC cells were detected in blood, lungs and liver of mice carrying MFP tumors without significant brain metastatic outgrowth, even in the presence of large >4 cm^3^ primary tumors (**Suppl. Fig. 2A-C**). Analysis of brain immune cells from mice carrying similar size primary tumors (**Fig 1B, C**) revealed that in the pre-metastatic brain niche, there already exist differences in the proportions of immune cells between E2 and E2-depleted mice. Brain myeloid cells (CD45⁺CD11b⁺) were the most abundant cell population encountered in all groups, with two distinct subsets based on CD45 expression levels: CD45^Intermedia (int)^CD11b^+^ cells, previously defined as microglia and CNS-macrophages^36^(MG/CNS MФ), and CD45^High(HI)^CD11b^+^, marking infiltrating bone marrow-derived macrophages (BMDM) (**Fig 1D-G**). MG/CNS MФ constituted the predominant immune cell population across all experimental cohorts (**Fig 1D**). While most of these cells displayed a CD16/32⁺ phenotype with no significant inter-group variability, E2-treated mice exhibited a significantly higher proportion of CD206⁺ cells compared to the letrozole-treated group (**Fig 1E**). In contrast, the infiltration of BMDM was significantly reduced in E2-treated mice compared to the OVX and OVX+ letrozole groups (**Fig 1F**). While CD206 expression remained comparable across all cohorts, BMDM in the OVX and OVX+ letrozole groups exhibited a marked increase in CD16/32⁺ expression (**Fig 1G**). T cell infiltration into the brain was significantly elevated in letrozole-treated mice compared to the E2-treated cohort (**Fig 1H**). This recruitment was characterized by a specific increase in the frequency of CD8⁺ T cells, whereas CD4⁺ T cell frequencies remained stable across groups (**Fig 1I, J**). Interestingly, within the total CD3⁺ T cell population, the proportion of CD8⁺ cells was unchanged, while the CD4⁺ subset was significantly reduced in letrozole-treated mice (**Fig 1K**). Collectively, these data indicate that aromatase inhibition promotes a preferential recruitment of cytotoxic T lymphocytes and repression of CD4⁺ T cells to the CNS in the pre-metastatic niche. We confirmed that this modulation of brain immune cells was not the result of exogenous luciferase or GFP expression in EO771 cells. In mice bearing MFP tumors from unlabeled cells, E2 treatment consistently reduced BMDM recruitment and suppressed CD16/32 expression, whereas letrozole treatment upregulated CD16/32 on these cells. Furthermore, the letrozole-induced expansion of CD8⁺ T cells and concomitant reduction in CD4⁺ T cells were maintained, resulting in a significantly elevated CD4/CD8 ratio in E2-treated mice compared to letrozole cohorts. These findings verify that the observed immunomodulatory effects are driven by hormonal status rather than reporter gene expression (**Suppl. Figure 3**). Notably, while letrozole showed the most robust immune shifts relative to E2, no significant differences were observed between OVX and OVX+ letrozole cohorts, suggesting low levels of peripheral (and brain E2) in the OVX mice are insufficient to modulate immune cell recruitment. Thus, the pre-metastatic brain niche of E2-treated mice presents a permissive niche for disseminated tumor cells characterized by suppressive anti-tumor myeloid cells and limited T cell recruitment.

**Figure 1.**
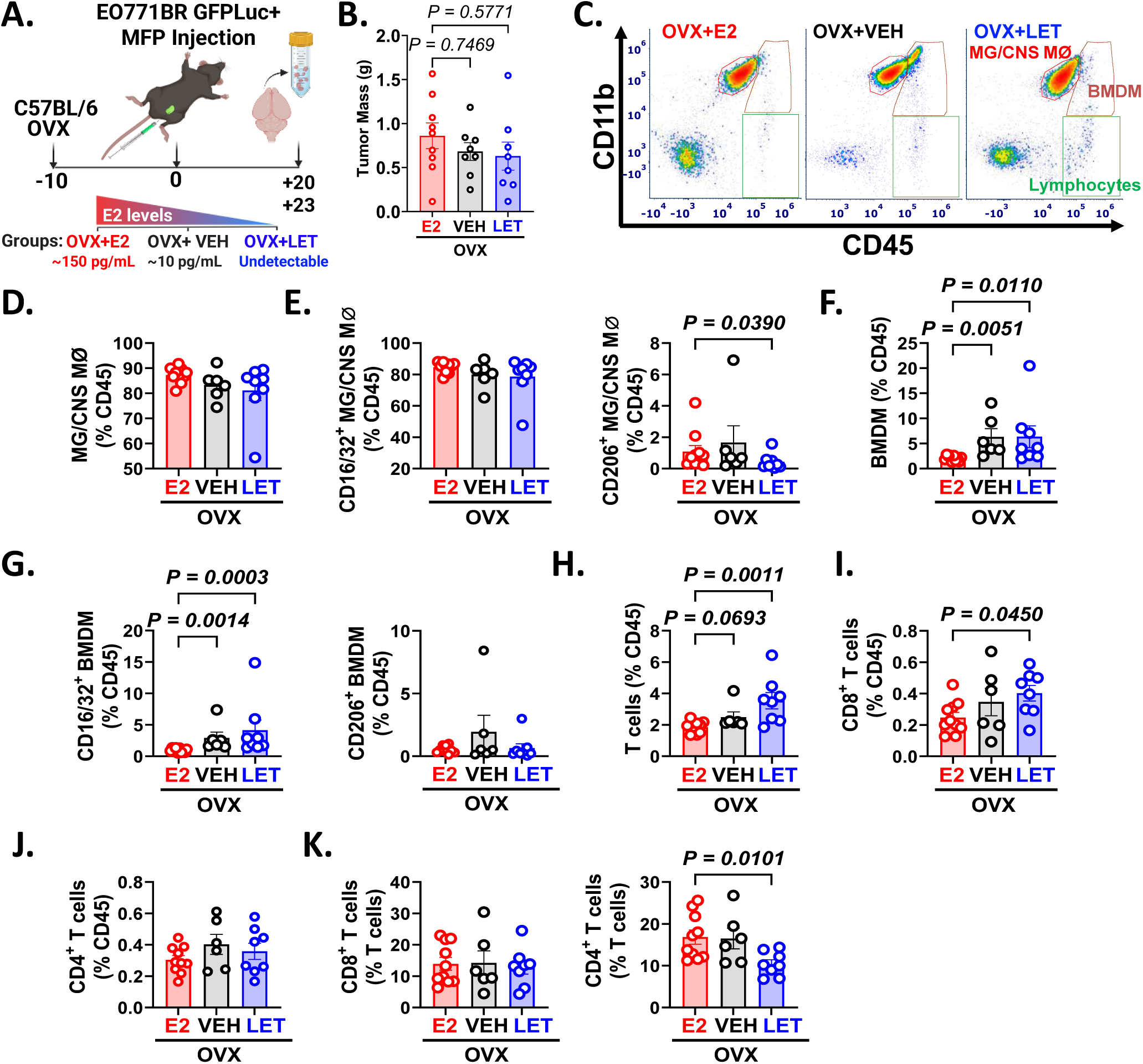
E2 represses effector-immune cell recruitment to the brain since early stages of spontaneous brain metastatic dissemination. **A.** OVX C57BL/6 were injected orthotopically with 1.2 ×10^6^ EO771BR1-GFPLuc+ cells in the MFP. Animals were randomized into three groups receiving (i) E2 pellet (OVX+E2, n=10), (ii) Vehicle (OVX, n=6) or (iii) letrozole (OVX+ LET, n=8) and euthanized when tumors reached similar size. Brain immune cells were isolated and analyzed using multiparametric flow cytometry. Colored label marks average serum E2 levels in each group. B. Graph shows mean (±SEM) of primary tumor mass per mouse at the time of euthanasia. Data was analyzed using one-way ANOVA. C. Representative image showing the major immune cell populations in the brains of mice across different treatment groups. D-K. Graphs show the percentage of indicated immune cell populations gated on CD45^+^ (D-J.) or CD3^+^ T (K.) cells per mouse (dots). Lines show the mean ±SEM. Pre-defined comparisons were analyzed using Kruskal-Wallis followed by uncorrected Dunn’s test. A. Created in https://BioRender.com. MG/CNS MØ=Microglia/ Central nervous system macrophages.

### Estrogen represses immune cells recruitment to the brain at later stages of brain metastatic progression

To assess how E2 modulates the brain immune microenvironment after cancer cells have colonized the brain, we analyzed the immune infiltrates in the brains of OVX C57BL/6 mice treated with E2, Veh, or letrozole, 12 days after IC injection of EO771BR1-GFPLuc+ cells. At this time point, E2-treated mice had significantly increased brain metastatic burden compared to OVX or OVX + Let-treated mice, as confirmed by *in vivo* and *ex vivo bioluminescence brain imaging* (**Fig 2A-C**). Immune profiling revealed that E2-treated mice showed an increased MG/CNS MФ fraction compared to the OVX+ Let-treated group; and while CD16/32⁺ expression remained unchanged, there was an increase in CD206⁺ expression within this population (**Fig 2D, E**). Conversely, E2 markedly reduced the infiltration of BMDM and suppressed their CD16/32⁺ pro-inflammatory polarization compared to both Veh and letrozole groups, without changes in the CD206^+^ levels (**Fig 2F, G**). Furthermore, E2-treated mice exhibited diminished CNS recruitment of total lymphocytes and CD8⁺ T cells relative to the letrozole cohort, while CD4⁺ T cell frequencies remained stable (**Fig 2H-J**). Notably, immune profiles were comparable between OVX and OVX+ letrozole mice, but letrozole-induced aromatase inhibition drives more robust immunomodulatory shifts than OVX alone, consistent with our observations in the pre-metastatic niche (**Fig 2H-J**).

**Figure 2.**
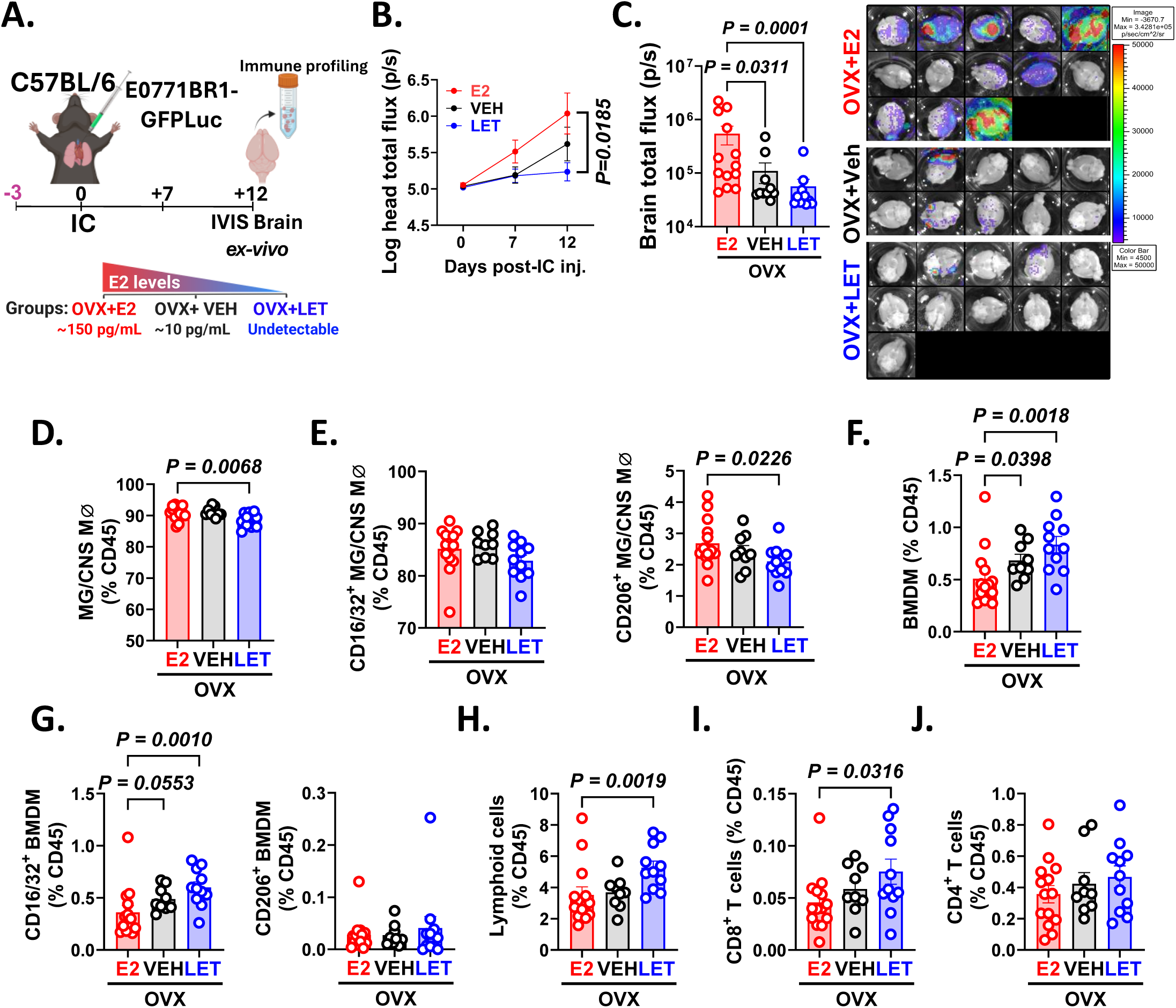
E2 induces a tumor-permissive immune environment during late-stage brain metastatic growth in the C57BL6/EO771 model. **A.** OVX C57BL/6 mice were randomized to receive (i) E2 pellet (n=13), (ii) Vehicle (n=10) and (iii) letrozole (n=11) three days prior to IC injection of 1×10^5^ EO771BR-GFPLuc+ cells and euthanized 12 days later. Brain immune cells were isolated and analyzed by multiparametric flow cytometry. **B.** *In vivo head* tumor progression followed by IVIS over time. **C.** Right panel: Brain *ex vivo* IVIS at euthanasia showing all brains analyzed. Left panel is quantification of right panel. **D-J.** Graphs show the percentage of indicated immune cell populations gated on CD45^+^ cells per mouse (dots); lines show the group mean ±SEM. Pre-defined comparisons were analyzed using one-way ANOVA followed by Fisher’s LSD test or Kruskal-Wallis followed by uncorrected Dunn’s test A. Created in https://BioRender.com. LET= Letrozole, MG/CNS MØ=Microglia/ Central nervous system macrophages.

To define whether similar changes existed across multiple models, we replicated the experiment in OVX-female BALB/c mice, supplemented with E2, Veh, or letrozole, injected with E2-unresponsive 4T1BR5 cells (**Suppl. Fig 4A**). Consistent with the EO771 model, E2-treatment significantly increased brain colonization compared to OVX and OVX+ Let-treated mice (**Suppl. Fig 4B**) and showed a reduced number of CD45⁺ brain immune cells (**Suppl. Fig 4C**). While microglia proportions were comparable across groups and primarily CD16/32⁺, E2 treatment specifically increased the CD206⁺ fraction and suppressed lymphoid cell infiltration and activated CD69^+^ CD8^+^ T cells (**Suppl. Fig 4D-H**).Thus, these results suggest that E2 promotes a tumor-permissive immune landscape from early through late stages of brain metastatic colonization in multiple E2-unresponsive BC-BM models.

### Estrogen promotes immunosuppressive transcriptional programs in ER^+^ microglia

Given that microglia, the most abundant immune cells in the brain, are primary mediators of E2 signaling via ER expression, we sought to characterize the E2-dependent transcriptional remodeling of the brain immune landscape. To this end, we performed scRNA-seq on brain cells isolated from OVX mice 10 days post-IC injection of EO771-GFPLuc+ cells following E2, Veh, or letrozole supplementation (**Fig 3A, Suppl. Fig 5A**). After filtering, droplet-based scRNA-seq resulted in 19990 total cells (**Suppl. Fig 5B**). Based on the expression of established marker genes and unsupervised cell type annotation, clusters were identified as microglia, dendritic cells, choroid plexus, fibroblast, macrophage, neutrophils, oligodendrocytes, NKT, B, astrocytes, pericyte, erythrocyte, and endothelial cells (**Fig 3B, C**). Consistent with our prior results, most of the profiled cells were microglia/dendritic cell clusters, without changes in the frequency of major cell types between treatment groups at this time-point (**Fig 3D, Suppl. Fig 6**). Since microglia was the most abundant immune cell population, we further clustered microglia into 6 subclusters using unsupervised clustering and overrepresentation analysis of their gene expression. Subclusters were identified as undifferentiated Microglia (Mg), proliferating Mg, interferon Mg, phagocytic Mg, migratory Mg and Tnf-alpha Mg (**Fig 3E-G**, **Suppl. Fig 7**). The migration Mg cluster, marked by microglial homeostatic genes known to promote activation and surveillance movements (*P2ry12, Sparc*)^37,38^, the TNF-α cluster, marked by genes associated with promotion of microglial activation (*Egr1, Fos, Nfkb1a*)^39,40^, and the Interferon Mg cluster, marked by genes involved in the regulation of type I Interferons (*Irf7, Ifit2/3, stat1*) were less abundant in E2-treated mice than OVX or letrozole-treated mice (**Fig 3G, H**). An undifferentiated Mg cluster, marked by cytoplasmic translation and translational genes (*Rsp29*, *Prl38*) was more abundant in E2-treated mice compared to OVX or letrozole-treated mice, while less abundant phagocytic and proliferating clusters were similarly represented among the 3 treatment groups (**Fig 3G**). Consistently, overrepresentation analysis showed microglia from E2-treated mice enriched in translation programs, while OVX (Veh) and OVX+ letrozole-treated mice had a significant enrichment of immune leukocyte activation myeloid cell migration and TNF signaling (**Fig 3H**). Microglia from OVX and OVX+ letrozole showed similar gene enrichment profiles, however letrozole was significantly enriched in hallmark IFN response and antigen processing programs (**Fig 3H**). Thus, these findings suggest that E2 treatment promotes a less activated, more homeostatic microglial phenotype compared to OVX or OVX+ letrozole treatment, as evidenced by the shift toward undifferentiated microglia and away from inflammatory subtypes.

**Figure 3.**
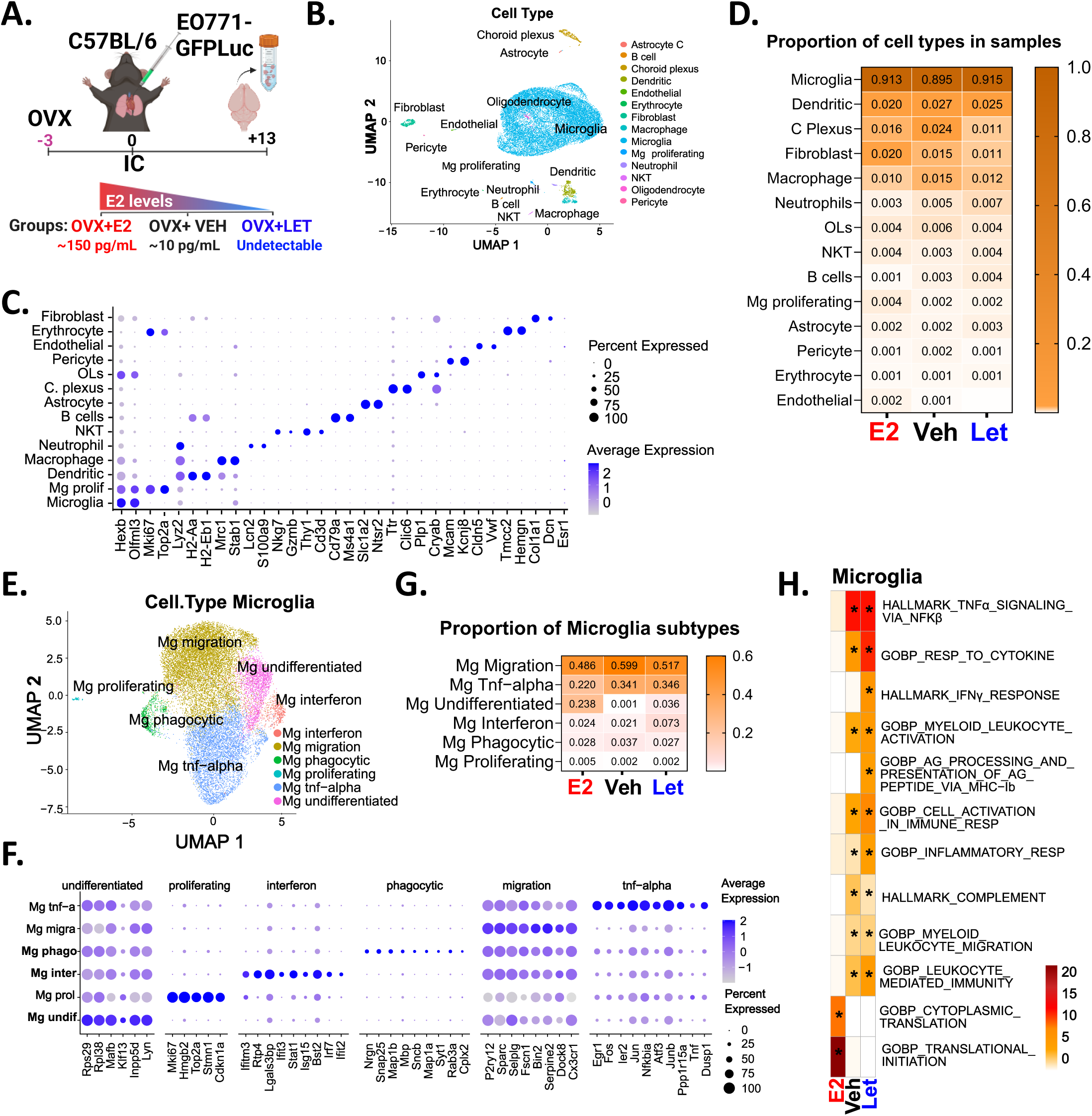
E2 suppresses immunosurveillance pathways in microglia/CNS-macrophages. **A.** OVX C57BL/6 mice received (i) E2 pellet (n=2 pooled), (ii) Vehicle (n=2 pooled) and (iii) letrozole (n=2 pooled) three days prior to IC injection of 1×10^5^ EO771BR-GFPLuc+ cells and euthanized 13 days later. Brain immune cells were isolated and analyzed by droplet based scRNA-seq. **B.** UMAP of cell types **C.** Dot plot of markers identifying cell types. **D.** Proportion of cell types by sample. **E.** UMAP of microglia subtypes, **F.** Dot plot of enriched markers by microglia subtype in E. **G.** Proportion of microglia subtypes by sample. **H.** Gene set enrichment analysis. Mg=Microglia. A. Created in https://BioRender.com. LET= Letrozole

### Depletion of microglia/macrophages reduces E2- associated brain metastatic progression

Following our observation that E2 induces an immunosuppressive microglial landscape while letrozole enhances immune activation, we utilized a depletion strategy to assess the necessity of the myeloid compartment in driving these divergent metastatic outcomes. We evaluated whether the absence of microglia and macrophages would offset both the pro-metastatic effects of E2 and the protective effects of letrozole. For this, E2- and letrozole-treated OVX female C57BL/6 mice were pretreated with control or chow containing 1 g/kg of the selective colony-stimulating factor 1 receptor (CSF1R) inhibitor PLX5622^41^ and injected intracardially with EO771BR1-GFPLuc+ cells **(Fig 4A**). Pharmacological depletion of microglia with PLX5622 appeared to attenuate E2-induced BM compared to E2-treated controls (**Fig 4B, C**), however, this combination was associated with reduced tolerability and early mortality in several mice. In the OVX+ letrozole cohort, CSF1R inhibition did not significantly further reduce *ex vivo* metastatic burden, though a non-significant trend toward slower longitudinal progression of the intracranial signal was observed (**Fig 4B, C**). We confirmed that PLX5622 effectively eliminated MG/CNS MФ in both E2- and letrozole-treated mice without altering total lymphoid recruitment (**Fig 4D, E**). Notably, microglial depletion significantly shifted the T cell profile, increasing the frequency of CD8⁺ T cells and activated CD69⁺ CD8⁺ T cells in both treatment groups, while reducing activated CD4⁺ T cells in letrozole-treated mice (**Fig 4F-I**). This resulted in a decreased CD4/CD8 ratio across both cohorts (**Fig 4J**), suggesting that microglia depletion reprograms the brain immune microenvironment toward a more cytotoxic, anti-tumoral state.

**Figure 4.**
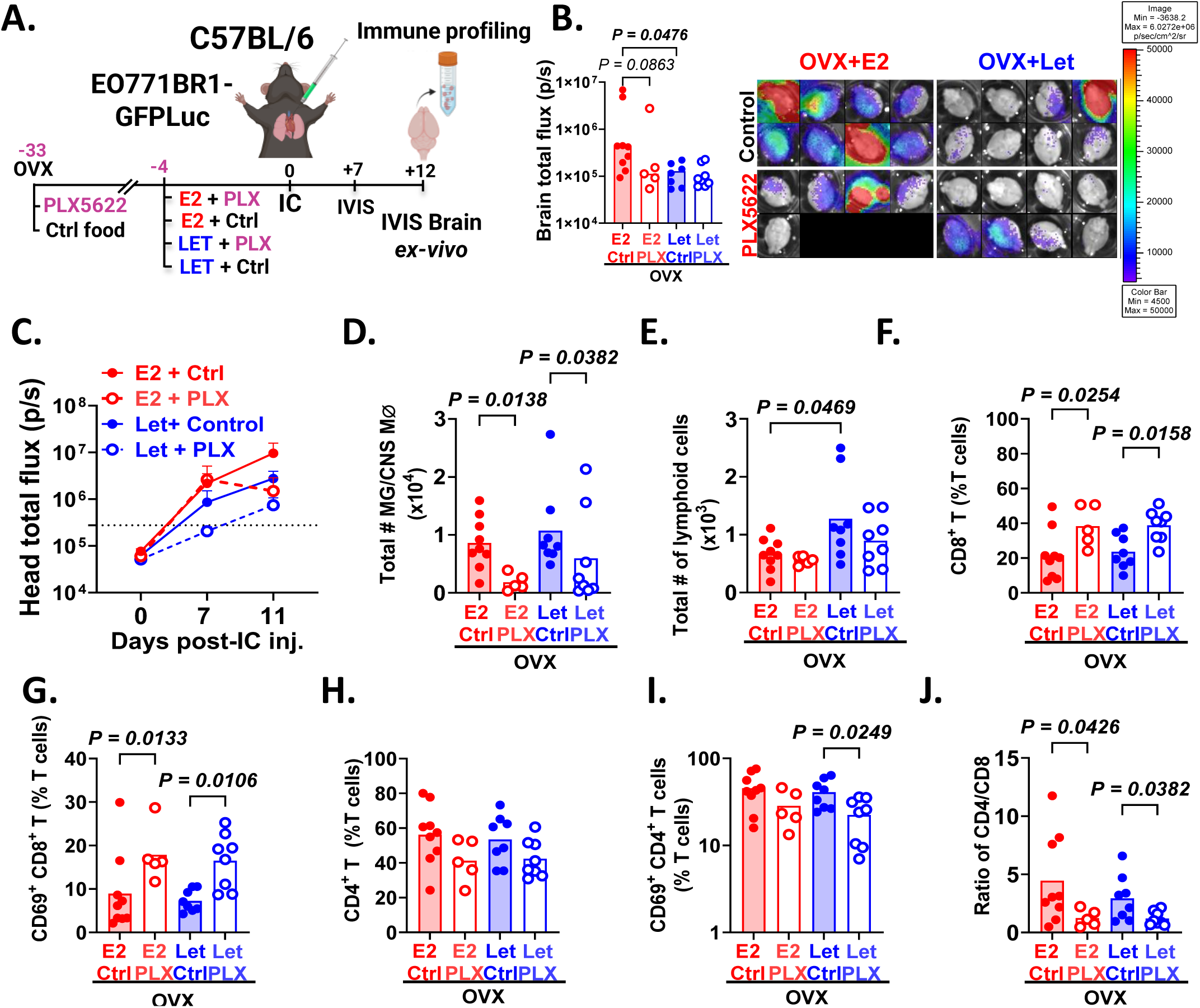
Microglia/macrophage depletion reduces E2-induced metastatic burden and promotes CD8^+^ T cell recruitment to the brain. **A.** OVX C57BL/6 mice were fed with PLX5622 or control chow for at least 15 days and then were randomized to the indicated treatments (n=5-9/group) four days prior to IC injection of 1×10^5^ EO771BR-GFPLuc. **B.** Left panel shows median of brain *ex vivo* IVIS quantification at euthanasia. Right panel shows image of all analyzed brains. **C.** Graph shows mean of head metastatic burden over time measured by IVIS. **D-E.** Total number of indicated immune cell type in each group. **F-I.** Graphs show the percentage of indicated immune cell populations gated on T cells per mouse (dots). **J.** Ratio of CD4^+^/CD8^+^ T cells. Pre-defined comparisons were analyzed using one-way ANOVA followed by Fisher’s LSD test or Kruskal-Wallis followed by uncorrected Dunn’s test. A graphic created in https://BioRender.com. LET= Letrozole, MG/CNS MØ=Microglia/ Central nervous system macrophages.

### Microglia from E2-treated mice show decreased ability to support T cell expansion and activation

Since E2 actively suppressed microglial differentiation toward inflammatory phenotypes and maintains a more quiescent microglial population, we next sought to determine how E2-mediated microglia reprogramming influences their ability to modulate T cell function using *in vitro* co-culture assays. To assess activation and effector potential, naïve T cells (isolated from non-tumor-bearing mice) were stimulated with αCD3/CD28 and co-cultured for 48 h with microglia harvested from BM-bearing ovariectomized (OVX-MG) or E2-treated (E2-MG) mice (**Fig 5A**). Both OVX-MG and E2-MG showed similar ability to increase activation markers (CD69) and granzyme B (GNZB) production in both CD4⁺ and CD8⁺ T cells compared with T cells cultured alone (**Fig 5B-E**). However, E2-MG significantly impaired the induction of IFN-γ in CD4^+^ and CD8^+^ T cell cells compared to OVX-MG (**Fig 5F, G**).

**Figure 5.**
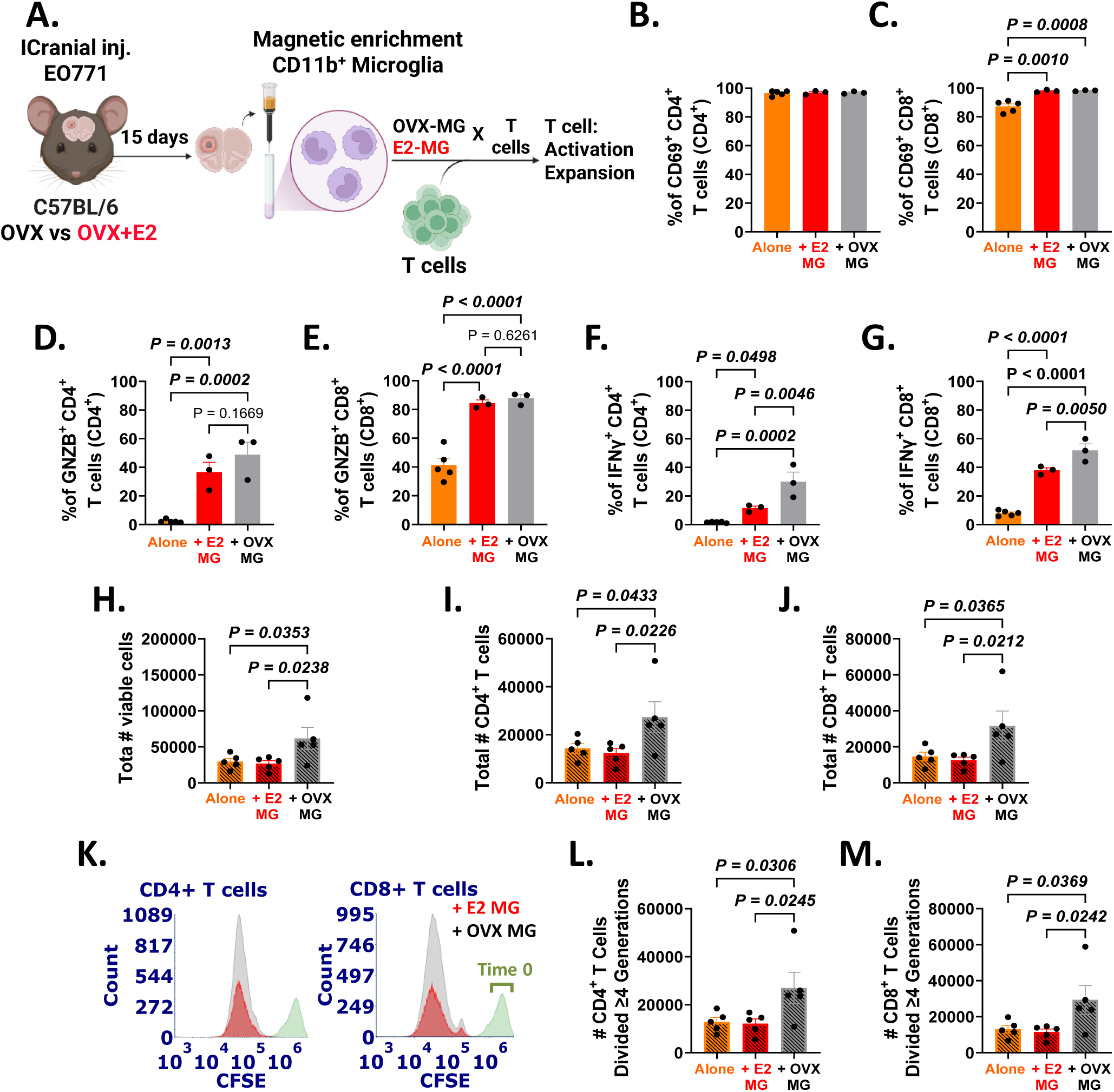
Microglia from E2-treated mice have reduced ability to support T cell expansion and activation. **A.** OVX C57BL/6 mice received either (i) E2 pellet or (ii) placebo pellet two weeks prior to intracranial injection of 5×10^4^ EO771 cells. Mice were euthanized 15 days later. CD11b⁺ microglia were isolated and cocultured with T cells from naïve C57BL/6 female mice to assess T cell activation and expansion. **B-G** T cell activation assays from microglia-T cell co-culture. Graphs show percentage of the indicated immune cell populations gated on CD4⁺ T or CD8⁺ T cells per replicate (dots). Bar indicates mean ± SEM. **I-M** T cell expansion assays from microglia-T cell co-cultures. **H-J.** Total number of the indicated cell populations per replicate (dots) after 48 h of co-culture. Bars are mean ± SEM. **K.** Representative histograms showing CFSE intensity and cell counts of CD4⁺ T cells (left) and CD8⁺ T cells (right) after two days of coculture with E2 MG or OVX-MG. Green histogram (day 0, T cells alone) indicates CFSE intensity of CD4⁺ or CD8⁺ T cells prior to co-culture**. L-M**. Total number of CD4^+^ and CD8^+^ T cells divided over 4 generations after 48 h of co-culture. Graphs show mean ± SEM. Pre-defined comparisons were analyzed using one-way ANOVA followed by Fisher’s LSD test. ICranial= Intracranial. A. Created in https://BioRender.com.

To evaluate T cell expansion, pre-activated T cells were labeled with CFSE and sub-cultured with microglia in the presence of IL-2. While OVX-MG robustly supported T cell expansion, E2-MG exhibited a diminished capacity to promote proliferation (**Fig 5H-J**). CFSE dilution analysis confirmed that OVX-MG significantly increased the number of cells undergoing at least 4 division cycles compared to E2-MG (**Fig 5K-M**). Collectively, these data indicate that E2 polarizes microglia toward an immunosuppressive phenotype that limits IFN-γ production and IL-2-driven mitogenic expansion, whereas E2 depletion (OVX) shifts microglia toward a state that supports anti-tumoral T cell responses.

### E2 suppression increases the effectiveness of brain radiation to decrease BM

Given the potential for E2-suppresion to reactivate anti-tumoral immunity in BM, we evaluated whether E2 depletion enhances therapeutic efficacy of WBRT. Since RT is part of the clinical standard of care for BM and potent modulator of antigen presentation; we specifically determined whether E2 suppression synergizes with RT to bolster anti-tumoral responses in preclinical models of late-stage BM. To mimic pre-menopausal BM formation, OVX C57BL/6 mice were supplemented with E2 pellets and injected intracardially with EO771BR1-GFPLuc+ cells. Seven days post-injection, once metastases were established, mice were randomized to continue E2, undergo E2-withdrawal (EWD), or receive EWD + letrozole (EWD+ Let), each with or without 15 Gy WBRT (**Fig 6A**).

**Figure 6.**
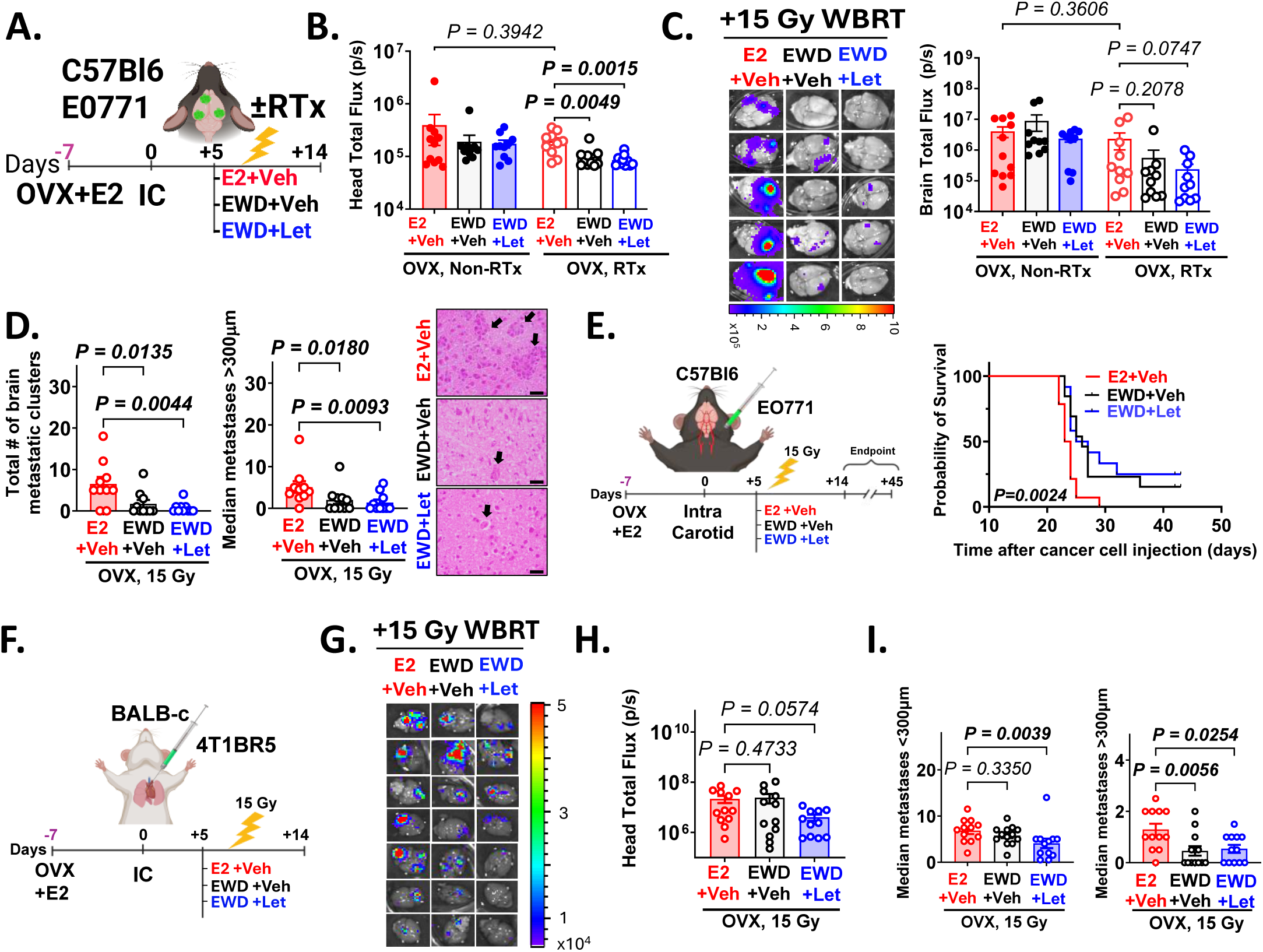
E2 suppression combined with WBRT decreases BM progression in immunocompetent mouse models of ERࢤ BC. **A.** OVX C57BL/6 mice receiving E2 were injected with 5×10^4^ EO771BR1-GFPLuc+ cells intracardially, and 5 days later were randomized into (i) E2, (ii) EWD, and (iii) EWD + Let (n=10/group). E2 and EWD mice received injections with vehicle control. Mice received a single dose of 15 Gy of WBRT 8 days post-injection and were euthanized 14 days later. **B.** Metastatic burden in the head measured right before euthanasia via IVIS in mice from A. **C.** Left: representative *ex-vivo* brain IVIS images. Right: median of brain total flux (photon/second) per mouse from A. **D.** Total number of brain metastatic clusters per mouse (left) and median number of metastases (>300 µm) per mouse (right). Representative H&E stain of brain metastatic clusters from mice is shown. **E.** OVX C57BL/6 mice receiving E2 were injected with 1×10^4^ EO771BR1GFPLuc+ cells via intracarotid artery injection. BM were allowed to grow for 5 days, and mice were randomized to (i) E2 (n=13), (ii) EWD (n=13), and (iii) EWD + Let (n=12). All mice received a single dose of 15 Gy of WBRT 8 days post-injection and were euthanized at human endpoints (Left). Graph shows Kaplan-Meier survival curve analyzed using a Log-rank test (Right). **F.** OVX BALB-c mice receiving E2 were injected with 2.5×10^4^ 4T1BR5-mCherryLuc cells via IC injection. BM were allowed to grow for 5 days, and mice were randomized into (i) E2, (ii) EWD, and (iii) EWD + LET (n=12 per group). All mice received a single 15 Gy WBRT dose 8 days post-injection and were euthanized 14 days later. **G.** Representative *ex-vivo* brain IVIS images from F. **H.** Head metastatic burden at endpoint from F. **I.** Histological quantification of metastases from F. Graphs show median number of micrometastases (<300 µm, left) and macrometastases (>300 µm) per mouse. For all graphs, data analysis was Kruskal-Wallis followed by uncorrected Dunn’s test. A, E, F. Created in https://BioRender.com. LET= Letrozole.

In contrast to the efficacy of OVX+ letrozole in *<u>preventing</u>* colonization (**Fig 2**), E2 suppression alone did not reduce established late-stage BM. However, combining WBRT with EWD or EWD+ Let significantly attenuated metastatic progression compared to E2-treated controls, as demonstrated by reduced head IVIS signal, *ex vivo* brain bioluminescence, and histological quantification (**Fig 6B-D**). To evaluate the survival impact of this synergy, mice bearing EO771BR1 intracarotid-induced brain-only metastases were treated with 15 Gy WBRT and randomized to E2, EWD, or EWD+ Let cohorts (**Fig 6E**). E2 suppression significantly extended survival from a median of 23 days (E2-only) to 26 days (EWD and EWD+ Let; *P=0.0024*), with 25% of mice exhibiting a doubling of overall survival (**Fig 6E**).

These findings were validated in the 4T1BR5 BALB/c model, where WBRT+EWD+ Let significantly reduced both head IVIS signal and histological tumor burden (**Fig 6F-I**). Notably, this combination therapy also reduced extracranial burden in the EO771/C57BL6 model (**Suppl. Fig 8A**) but not in the 4T1BR-BALB/c model (**Suppl. Fig 8B**), suggesting a cell-line or host-specific systemic anti-tumor mechanism. To confirm that this therapeutic response requires functional T cell recruitment, we repeated these studies in severely immunocompromised NSG mice. In the absence of an intact immune system, EO771BR1 brain and systemic metastases were entirely refractory to EWD, regardless of WBRT status (**Suppl. Fig 8C-D**). Collectively, these results demonstrate that E2 suppression leverages microglial modulation to reactivate anti-tumoral T cell responses, thereby enhancing the efficacy of radiation therapy in late-stage BM.

## DISCUSSION

It is now recognized that estrogens play pro-tumorigenic roles that function beyond their well-known mitogenic activity in ER^+^ tumor cells. Here we show for the first time that E2 reprograms ER^+^ microglia/CNS-macrophages and modulates the recruitment of anti-tumoral peripheral effector immune cells to the brain to promote BC-BM. Our data shows that reprogramming of microglia/CNS-macrophages and increased immune T cell infiltration to the brain occur before the establishment of clinically detectable metastases in spontaneous BM models, indicating that systemic estrogen levels can “prime” the brain environment to favor the survival of disseminated tumor cells before they even colonize the tissue.

Our scRNA-seq analysis of brain immune cells showed microglia as the most abundant immune cell in BM from E2 or E2-suppressed mice. Microglia have been shown to promote anti-tumor natural killer and T cell responses and suppress BM^23^, or play an immunosuppressive pro-tumorigenic role at later stages of BM progression^22^, but whether E2 impacts microglia function in the context of BM is less known. In contrast to prior reports indicating a role for E2 in modulating macrophage and microglia M1/M2 polarization^42^, our scRNA-seq and immune profiling studies demonstrate that E2 does not promote clear M1/M2 polarization of microglia in BM, but rather a more complex shift in their transcriptomic states. Collectively, our data demonstrates that E2 enforces a transcriptional state of functional quiescence in microglia, characterized by an expansion of undifferentiated populations and the suppression of activation programs. While E2 deprivation (OVX and letrozole) shifts the microglial landscape toward pro-inflammatory signaling (TNFα, IFN), surveillance (migratory), and antigen-presentation, E2 maintains a more immature and translationally focused phenotype. Notably, letrozole treatment provides a more robust activation of interferon-responsive genes than ovariectomy alone, highlighting its potency in reversing E2-mediated immune suppression within the brain.

Our studies showed that E2 suppressed the fraction of CD8^+^ T cells to the brain while maintaining CD4^+^ T cell frequencies, resulting in an elevated CD4/CD8 ratio. This pattern persisted from the pre-metastatic niche through established brain metastases and was reversed by letrozole treatment, which preferentially increased CD8^+^ T cell infiltration. This repression of cytotoxic CD8^+^ T cells may result from repression of migratory and IFN clusters in microglia, which are typically responsible for the chemotactic signals required for T cells recruitment^43^, as well as the decreased ability to promote CD8^+^ T cell expansion. The fact that microglia depletion did not alter the ability of E2 or letrozole to recruit T cells to the brain, yet it significantly enhanced the percentage of CD8^+^ T cells, suggest direct influence of microglia to modulate the functionality of T cells. In contrast to prior studies where E2 stimulated macrophages suppressed CD8^+^ but not CD4^+^ T cells^27^, our results showed that E2 reduces the ability of microglia to promote expansion of both CD8^+^ and CD4^+^ T cells, suggesting brain-associated microglia/macrophages may have additional immunomodulatory functions than peripheral macrophages.

Our study has some limitations. While gene signatures used to assign cell types in scRNA-seq identify mostly microglia clusters and a small population of macrophages in the brain^44^, the specific contribution of E2 to transcriptional reprogramming of microglia vs macrophages remains unknown. Immuno-labeling of activated microglia/CNS-macrophages (CD45^int^/CD11b^+^) and BMDM (CD45^HI^/CD11b^+^)^36^ used in our studies suggests that E2 differentially regulates activation and recruitment of resident CNS microglia and macrophages to the brain. Since macrophages and resident microglia have been shown to play a different role from macrophages in the promotion of brain tumors^45,46^, future studies are warranted to dissect the specific contribution of microglia and BMDM subpopulations to endocrine therapy and WBRT responses.

Importantly, the observation that OVX (which results in low levels of peripheral E2) and OVX+ letrozole (which results in complete E2-depletion) have different abilities to block myeloid cell activation and recruitment to the brain, suggests that E2/ER dynamics influenced by diverse endocrine therapies may differentially alter microglia vs myeloid cell function. Future studies are needed to determine if other endocrine therapies, namely selective-ER degraders (SERDs) or selective ER-modulators (SERMs), have similar effects on immune cell function in BM.

Finally, our studies have important therapeutic implications. Endocrine therapies are emerging as attractive therapeutic candidates in the multimodality treatment of metastases^47^. The consistent increase in T cell infiltration to the brains of E2-depleted mice in our studies would predict a TME able to effectively target BM, yet endocrine therapy alone was not sufficient to eliminate late BM but rather increased the effectiveness of brain radiation. It is possible that the synergism between E2 depletion and radiation results from the increased antigen release, secondary to radiation, plus the enhanced antigen presentation ability of microglia to T cells. Further studies are needed to specifically determine the extent to which antigen presentation capability plays a role in the antitumoral response in radiation and E2-suppressed mice. Nonetheless, these studies provide preclinical evidence that E2-suppression may have clinical utility in preventing or treating brain metastasis even in patients with ER-negative primary tumors by “re-awakening” the brain’s immune landscape.

## METHODS

### Cell lines

Murine E2-unresponsive EO771 cell line (RRID: CVCL_GR23) was purchased from CH3 BioSystems, LLC (Buffalo, NY), labeled with GFP-luciferase and passed through the mouse brain (EO771BR1) and cultured in RPMI-5% FBS. Murine brain trophic E2-unresponsive cell line 4T1 (4T1BR5, a gift from Dr. Suyun Huang) was modified to express mCherry-Luciferase and was cultured in DMEM/High Glucose with 5% FBS. For all *in vivo* experiments, cells were used within 5 passages from thawing. Cells were authenticated by STR testing at the UC-AMC Cell Technologies Shared Resource (RRID: SCR_021982) prior to cryopreservation and tested for mycoplasma (MycoAlert^TM^ PLUS-Lonza) every three months.

### Animal experiments

All animal experiments were approved by the University of Colorado Institutional Animal Care and Use Committee, and all animal interventions were performed by investigators blinded to the experimental groups. To model *brain metastatic colonization*, EO771, EO771BR1-GFPLuc, or 4T1BR5 mCherryLuc cells were injected via intracardiac (IC), intracarotid artery injections, or intracranially into 8-14-weeks old female C57BL/6J, BALB-c or NOD-SCID IL2Rγnull (NSG). To model *spontaneous BC dissemination*, mice received an injection of cancer cells in the fourth mammary fat pad (MFP). To model *pre-menopausal E2 levels,* mice were OVX and then supplemented with premenopausal E2 levels (required to promote growth of ER^+^ tumors in mice) via slow-release silastic pellets (0.5 or 1 mg), resulting in average of 150 pg/mL serum E2. To model E2 suppression, mice were OVX and treated subcutaneously (SQ) with vehicle (OVX, resulting in average 10 pg/mL serum E2) or 10 µg (50 µL) letrozole/daily (OVX+LET results in undetectable serum E2 levels). Letrozole stock was prepared at 0.2 mg/mL in 0.3% hydroxypropyl cellulose, and control mice received 50 µL vehicle. For microglia/macrophage depletion, PLX5622 (HY-114153, MedChemExpress) was incorporated into rodent diet AIN-76A (D10001) by Research Diets at 1g/Kg. Non-supplemented AIN-76A was used as the control diet. Mice were maintained on PLX5622-containing or control chow until the experimental endpoint (euthanasia).

### Brain radiation

Whole brain radiotherapy (WBRT) in mice was performed using the Precision X-Ray X-Rad 225Cx Micro IGRT and SmART Systems, using doses previously described to be non-lethal for immunocompromised mice (10 Gy), and for immunocompetent mice (15Gy)^28^ as indicated.

### *In vivo* luminescence (IVIS)

For *in vivo* analysis of metastases, mice received 200 µl of D-Luciferin, Potassium Salt (GoldBio LUCK-1G) [15 mg/mL] via intraperitoneal (IP) injection, and head and extracranial metastatic burden were calculated at the indicated times using Living Image® 4.8.2. Dissected brains were submerged into a Luciferin solution of 0.15 mg/mL in PBS for 10 min, and brain metastatic burden was measured via IVIS.

### Histological quantification of metastases

Micro- and macro-metastases were quantified histologically as previously described^16^. Briefly, six hematoxylin and eosin (H&E)-stained serial sections 300 µm apart in a sagittal plane through one brain hemisphere were analyzed at 4X magnification using an ocular grid, and the median number of metastases > and < 300 µm in each section was tabulated.

### Cell preparation for single-cell RNA sequencing (scRNA-seq)

Brain immune cells from adult mice carrying BM were isolated following the protocol originally outlined by Manglani *et al*^29^ with minor adjustments (**Suppl. methods**), and pooled from 2 mice per treatment. Viable cells were sorted using propidium iodine (PI) staining and 25,000-50,000 cells per sample were submitted to the Genomics and Shared Resource at UC-AMC (RRID: SCR_021984) for 10X Chromium capture.

### scRNA-seq processing and analysis

Cell Ranger (v4.0.0)^30^ was used to process the FASTQ files to cell and gene count tables using unique molecule identifiers (UMIs) aligning to the mouse genome mm10 (compiled by 10X Genomics as refdata-gex-mm10-2020-A). Downstream quality control and analysis utilized the Seurat (v4.3.0)^31^ pipeline. Cell Ranger-filtered data was further processed by removing genes identified in fewer than 10 cells. Cell barcodes were removed if they had greater than 15,000 UMIs, expressed fewer than 500 genes or greater than 4,000 or greater than 5% of UMIs from mitochondria. This filtering criteria resulted in 19,990 cells across all samples, totaling 18,132 genes. The gene counts were normalized using Seurat, where counts are divided by each cell’s total counts and multiplied by 10,000, followed by natural-log transformation. The top 2,000 most variable genes were scaled with total UMI and percentage mitochondria regressed out. Clusters were identified as cell types through multiple methods. The top 100 enriched genes by cluster were used as input for over-representation analysis^32^ with gene sets from the MSigDb C8 collection^33,34^ and the PanglaoDb^35^. Similar over-representation analysis was performed across microglia subtypes and across sample conditions using the Hallmarks and Gene Ontology biological processes collections. Canonical cell type markers and other genes were plotted in UMAP space and with dot plots using Seurat^31^. Microglia cells were subset, reclustered, and analyzed separately. Raw and processed single cell data are deposited in the Gene Expression Omnibus (GSE267950).

### Multiparametric flow cytometry staining and analysis

Brain immune cells were isolated as described above and stained (**Suppl. Methods**), using a panel of antibodies developed to examine the functional populations of immune cells, and activation, cytotoxicity, and expansion of T cells (**Suppl. tables 1-2)**. Flow cytometry was performed in a spectrum flow cytometer, 5-Laser Aurora system (Cytek Biosciences) using SpectroFlo 3.0, and FCS data were analyzed using FCS Express 7. The gating strategy used is shown in **Suppl. Fig. 9.**

### T cell-microglia co-culture assays

Primary T cells were isolated from spleen and pooled inguinal, brachial, and axillary lymph nodes using the Pan T Cell Isolation Kit II, mouse (Miltenyi Biotec, 130-095-130). Microglia were isolated from tumor-bearing brains using CD11b Microglia MicroBeads, human/mouse (Miltenyi Biotec, 130-093-634) and used immediately for co-culture experiments. T cells and microglia were cultured in phenol red-free RPMI-1640 supplemented with L-glutamine and containing 10% CS-FBS, sodium pyruvate, penicillin/streptomycin, gentamicin, β-mercaptoethanol, and nonessential amino acids. Mouse αCD3ε (5 µg/mL, InVivoMAb; Bio X Cell, BE0001-1) was used to coat plates overnight, and mouse αCD28 (1 µg/mL, InVivoMAb; Bio X Cell, BE0015-1) was added at the time of T cell seeding. For expansion assays, T cells were activated on αCD3-coated plates with soluble αCD28 for 48 h, stained with 0.5µM CFSE cell division tracker kit (Biolegend 423801), and subsequently re-plated in co-culture with microglia for an additional 48 h. Where indicated, 50 U/mL of recombinant mouse IL-2 (R&D Systems, 202-IL-010) was added. T cell proliferation and expansion were assessed by CFSE dilution and total cell counts by flow cytometry.

### Statistical Analysis

Data statistical analysis and graphs were completed using GraphPad Prism software version 10.1.0. Blinding and randomization were used for all mouse studies. Data normality was assessed by the Shapiro-Wilk and Kolmogorov-Sminov tests. Normal data was analyzed using one-way ANOVA or unpaired t-tests. Non-normally distributed total flux data was log-transformed [Y=log(Y)]. In cases where data was not normal after transformation, non-parametric methods (Kruskal-Wallis and Mann-Whitney U test) with multiple comparison tests were used as appropriate. A P value ≤0.05 was considered significant.

During the preparation of this work the author(s) used Co-pilot for language editing of introduction and abstract. After using this tool/service, the authors reviewed and edited the content as needed and take full responsibility for the content of the published article

## Ethics

All animal experiments were approved by the University of Colorado Institutional Animal Care and Use Committee.

## Data Availability

Raw files with proper identifiers for scRNA-seq can be accessed through GSE267950. Analyzed data will be made available upon reasonable request.

## Acknowledgments.

We thank the University of Colorado Cancer Center shared resources (Animal imaging and radiation *(RRID: SCR_021980),* Flow Cytometry *(RRID: SCR_022035),* Genomics *(RRID: SCR_021984)*, Biostatistics and Bioinformatics and the Human Immune Monitoring Shared Resource *(RRID:SCR_021985)* that are supported by NCI P30CA046934 and CTSA UL1TR001082 Center grants. Special thanks to Benjamin Van Court, Brooke Neupert and Tyler Weiskopf for management of the Small-Animal Irradiator Shared Resource.

