## Supplementary figures for "Estradiol Reprograms Microglia to Create an Immune-Suppressed Niche Permissive to Breast Cancer Brain Metastasis"

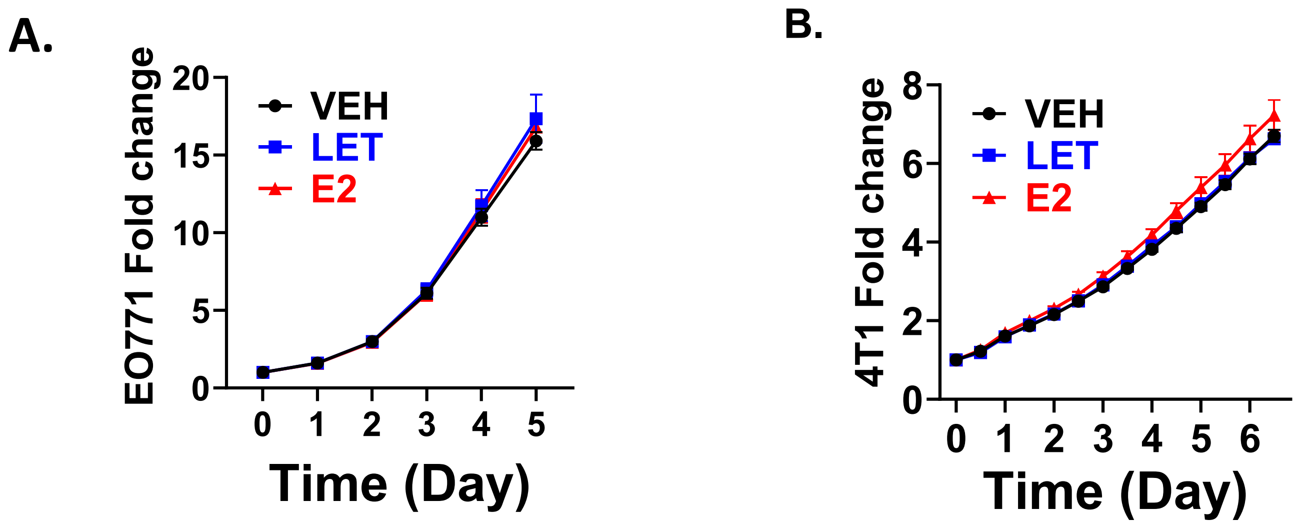


**Supplementary Figure 1.** EO771 (**A**) and 4T1 (**B**) cells treated with E2 (10nM), Letrozole (100nM) or vehicle (0.1% EtOH) and cultured in CS-FBS phenol red-free media. Graphs show fold change in confluence ±SEM relative to time 0, measured by live-imaging Incucyte. Data was analyzed using two-way ANOVA followed by a Fisher's LSD test. LET= Letrozole. There were no statistical differences at any time point.

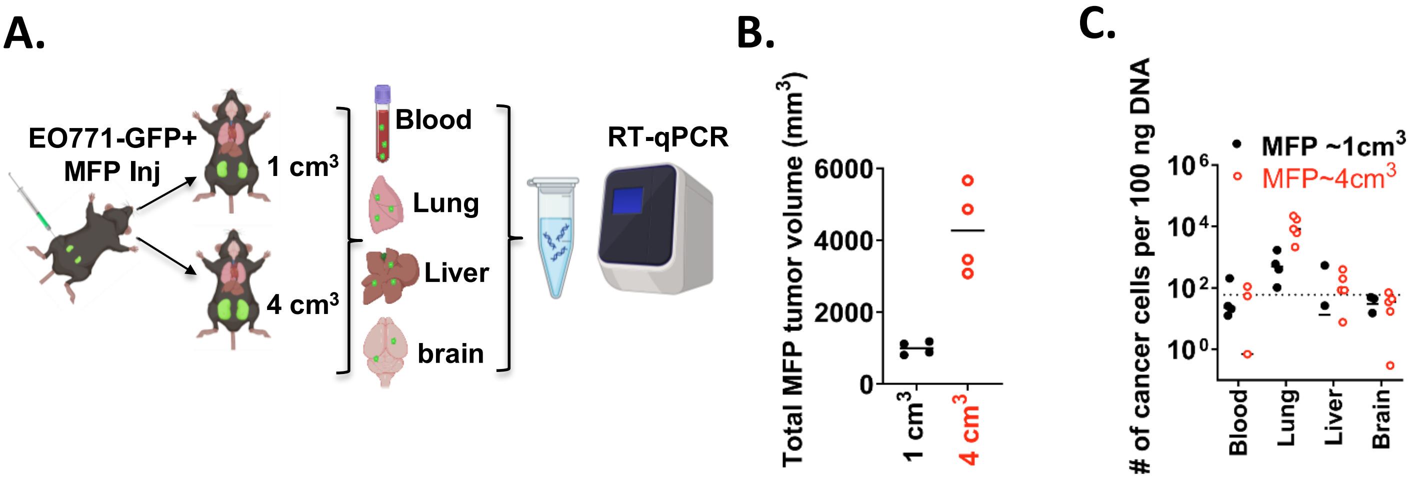


**Supplementary Figure 2. Detection of spontaneously disseminated cancer cells according to primary tumor size. A.** Female C57BL/6 were injected orthotopically with 3 x$10^6$ EO771BR-GFPLuc cells in the fourth MFP and 1 mg E2 pellet was implanted via SQ. Mice were euthanized when any of the tumors reached 1 $cm^3$(n=4) or 4 $cm^3$ (n=5). Blood was collected at euthanasia and mice transcardially perfused with cold PBS. Total DNA from blood and tissues indicated were used to quantify the number of eGFP+ cancer cells via RT-qPCR. **B.** Combined total tumor volume for mice in A. **C.** Number of disseminated eGFP+ cancer cells per 100ng DNA in samples from A, Dotted line indicates lower limit of detection. A. Created in https://BioRender.com.

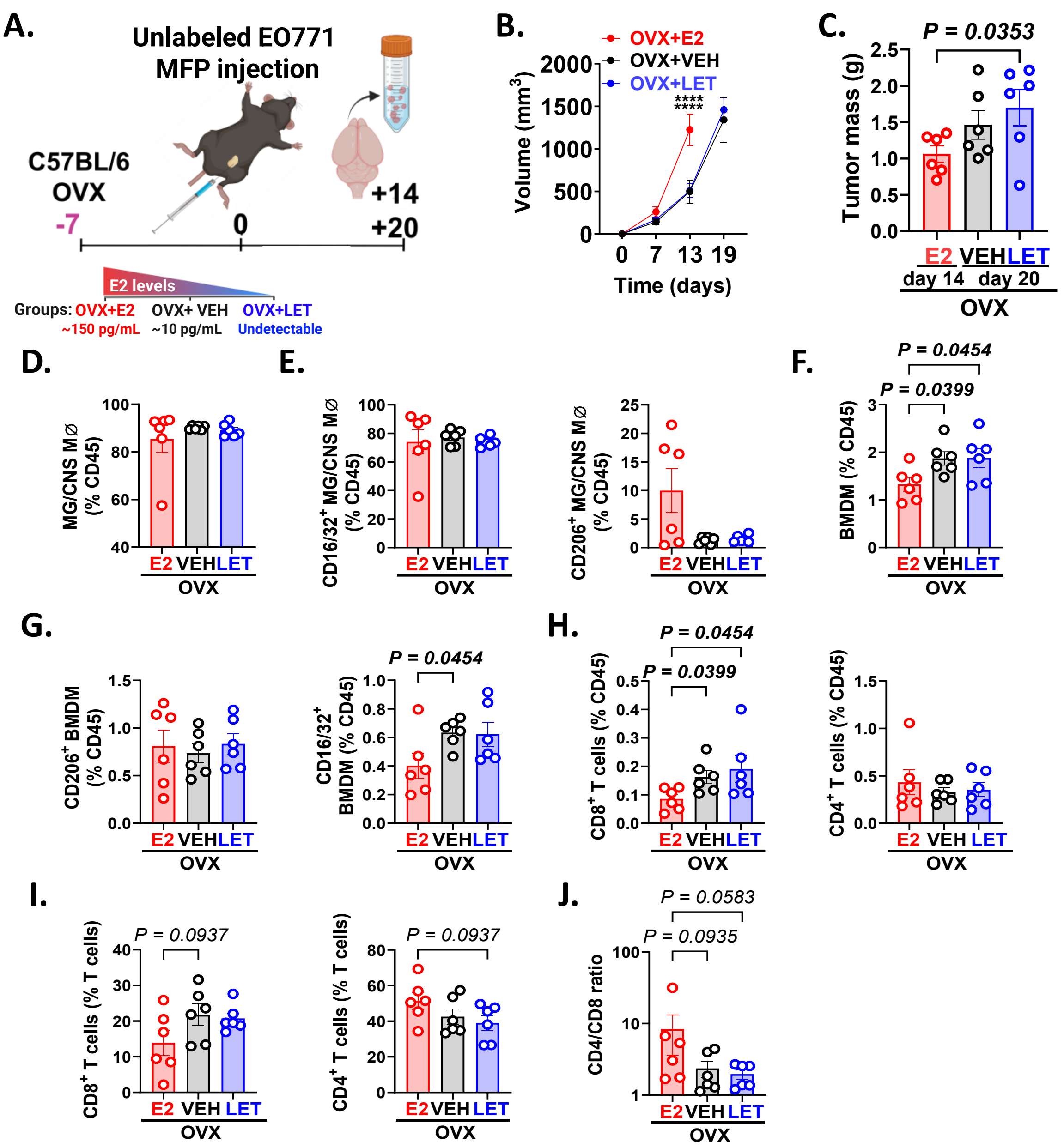


**Supplementary Figure 3. A.** OVX C57Bl/6 were injected orthotopically with 1.2 x$10^6$ EO771 cells in the MFP. Animals were randomized into three groups receiving (i) E2 pellet (OVX+E2, n=6), (ii) Vehicle (OVX, n=6) or (iii) letrozole (OVX+LET, n=6) and euthanized when tumors reached similar size (day 14 for E2-treated mice and day 20 for E2-depleted mice). Brain immune cells were isolated and analyzed using multiparametric flow cytometry. **B-C** Graphs show mean of primary tumor volume (B) and mass (C) ±SEM per mouse at the time of euthanasia (14 or 20 days). Data was analyzed using one-way ANOVA (C) and two-way ANOVA followed by Tukey correction (B). **D-J.** Graphs show the percentage of indicated immune cell populations gated on $CD45^+$ cells per mouse (dots). Lines show the mean ±SEM. Pre-defined comparisons were analyzed using Kruskal-Wallis followed by uncorrected Dunn's test. A. Created in https://BioRender.com. LET= Letrozole, MG/CNS MØ=Microglia/ Central nervous system macrophages.

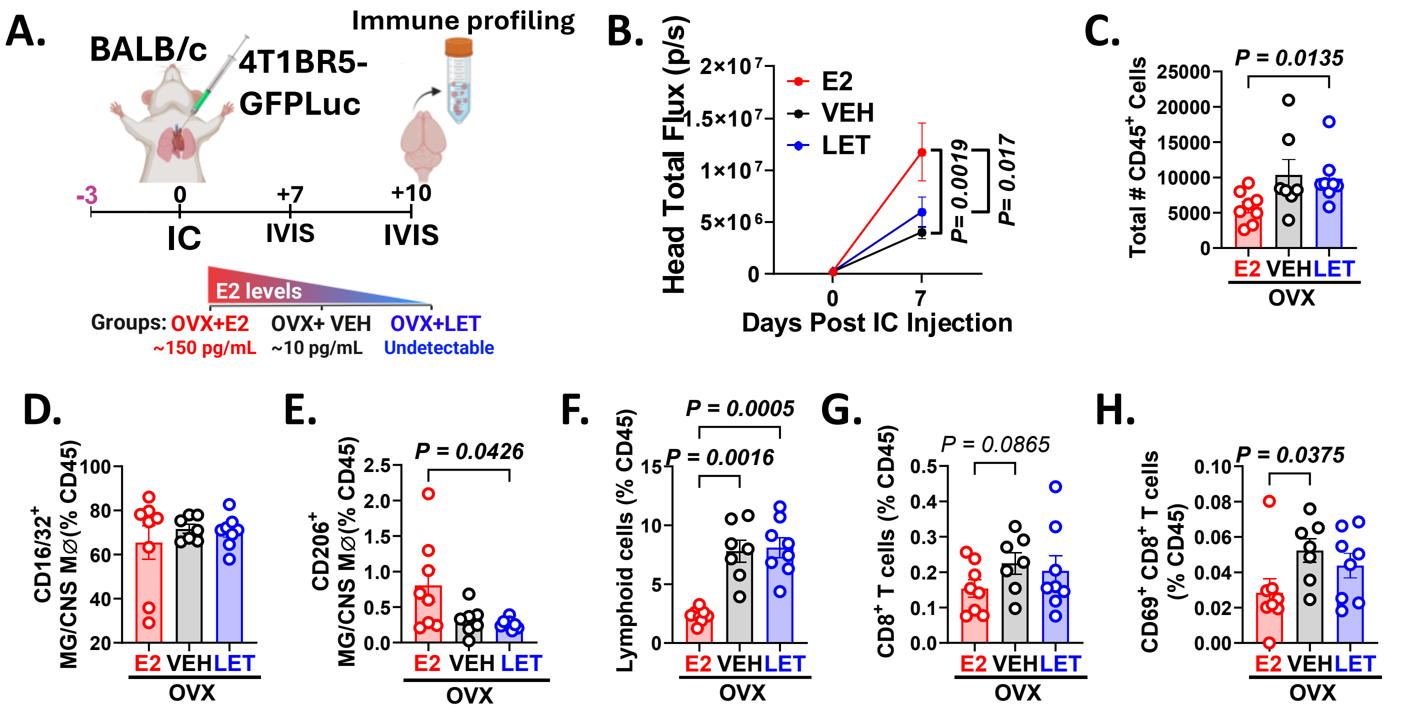


**Supplementary Figure 4. E2 induces a tumor-permissive immune environment in late stages of brain metastatic colonization in the BALBc-4T1BR5 model A.** OVX BALB-c mice were randomized to receive (i) E2 pellet (n=8), (ii) Vehicle (n=7) and (iii) letrozole (n=8) three days prior to IC injection of $2.5x10^4$ 4T1BR5-GFPLuc+ cells and euthanized 8 days later. Brain immune cells were isolated and analyzed by multiparametric flow cytometry. **B.** *In vivo* head tumor progression followed by IVIS over time. **C.** Total number of $CD45^+$ cells per group **D-H.** Graphs show the percentage of indicated immune cell populations gated on $CD45^+$ cells per mouse (dots); lines show the group mean ±SEM. Pre-defined comparisons were analyzed using one-way ANOVA followed by Fisher's LSD test or Kruskal-Wallis followed by uncorrected Dunn's test. A. Created in https://BioRender.com. LET= Letrozole, MG/CNS MØ=Microglia/ Central nervous system macrophages.

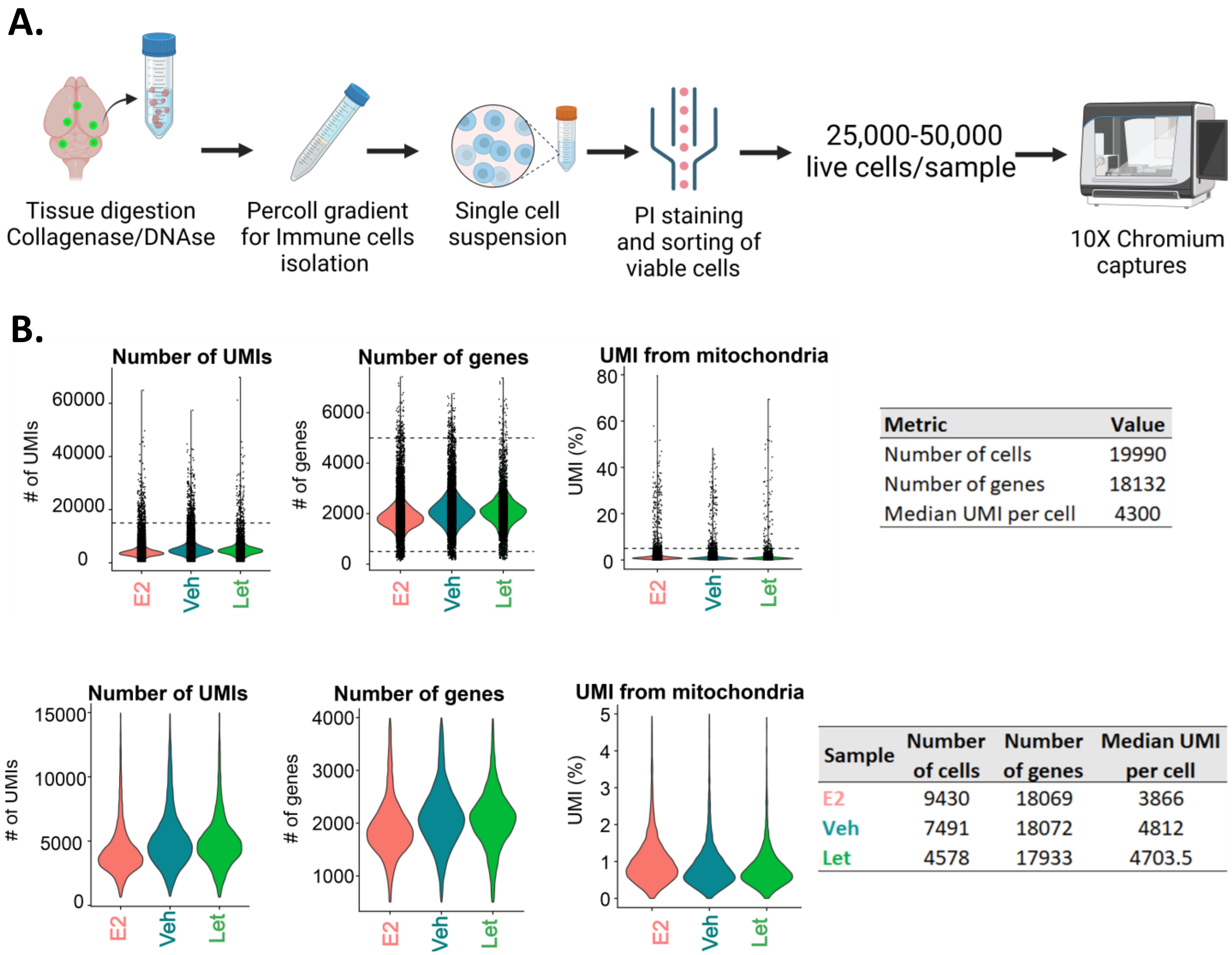


**Supplementary Figure 5. A.** Single-cell isolation pipeline, brain tissues were digested in 1 mg/mL collagenase D and 50 µg/mL DNAse in RPMI phenol red-free media, for 20 min, 37˚C. Brain immune cells were isolated using a percoll gradient (90%, 60% and 40%) and viable cells were sorted using propidium iodine (PI) staining. Between 25,000-50,000 cells per sample at 700-1100 cells/µl were submitted to the Genomics and Microarray Core at UC-AMC for 10X Chromium captures. **B.** Pre-filtering (Top) and post-filtering (bottom) quality control plots and tables. A. Created in [https://BioRender.com](https://BioRender.com).

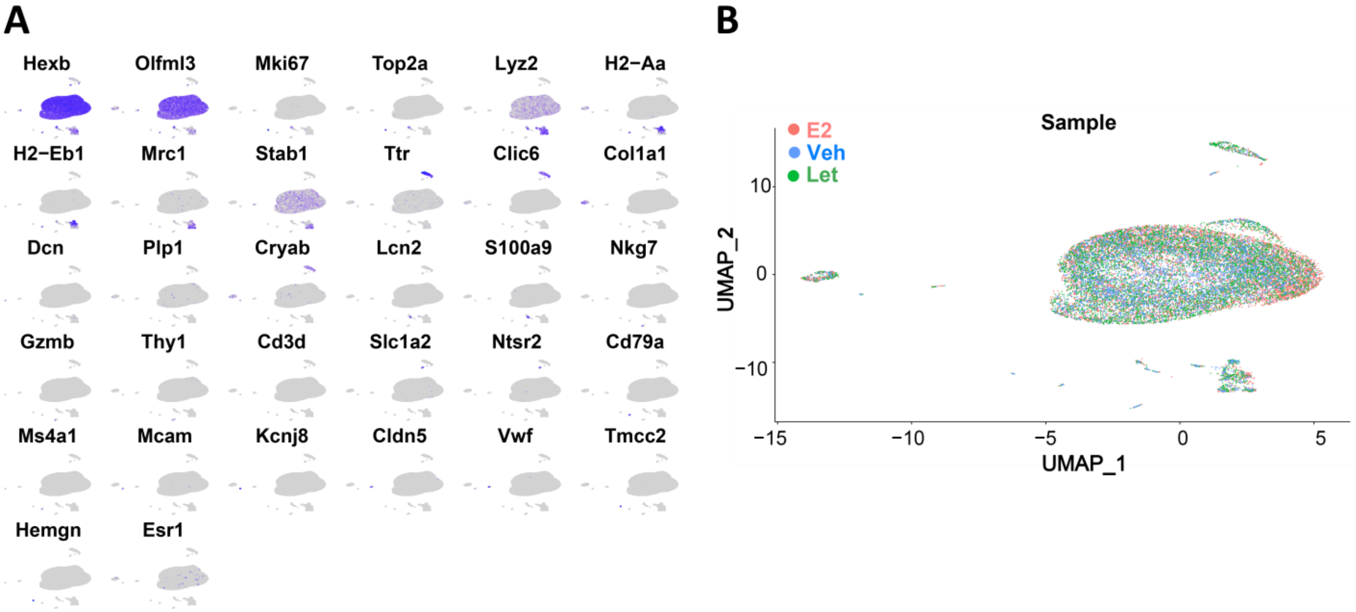


**Supplementary Figure 6. A.** UMAP cell-type marker grid sample supplementary to figure 3B. **B.** UMAP by sample supplementary to figure 3B.

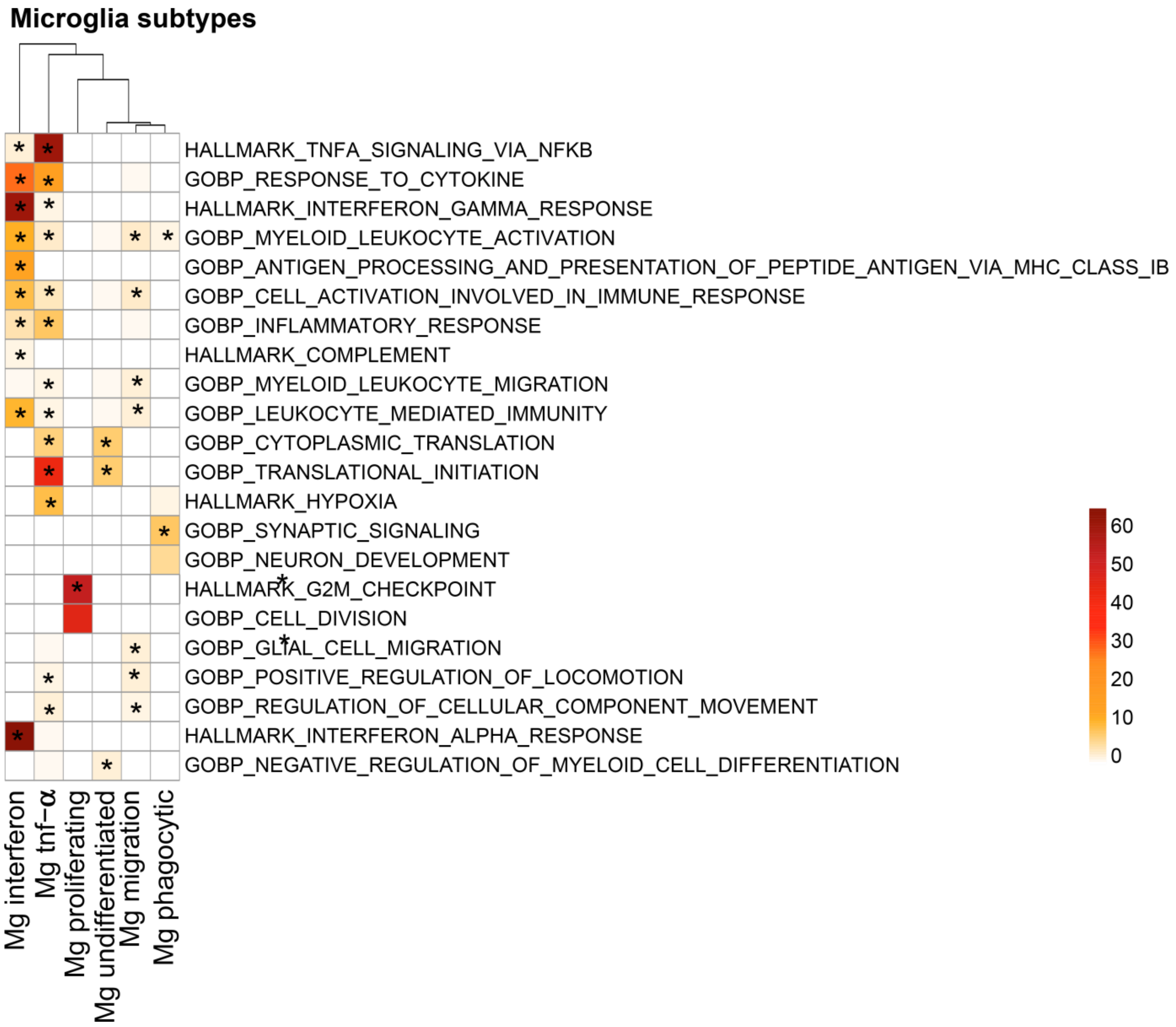


**Supplementary Figure 7.** Enriched gene sets in microglia subtypes supplementary to figure 3F.

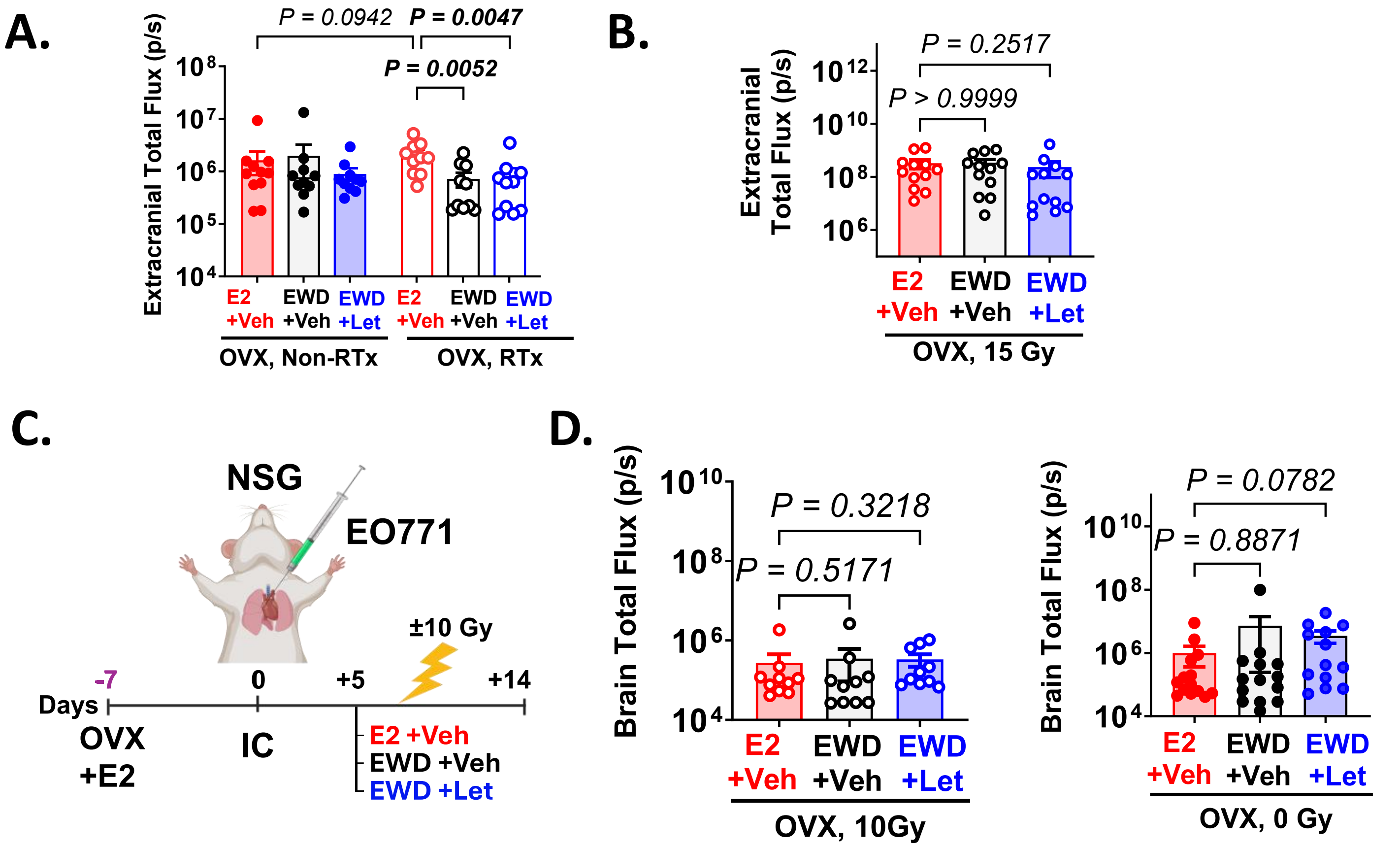


**Supplementary Figure 8. E2 suppression does not alter BM progression in immunocompetent models in the absence of brain radiation or in immunocompromised mouse models of ER$^-$ BC. A.** Extracranial tumor burden measured right before euthanasia via IVIS in C57BL/6 mice from fig 6A. **B.** Extracranial tumor burden measured right before euthanasia via IVIS in BALB-c mice from fig 6F. **C.** OVX NSG mice supplemented with E2 received 5x10$^4$ EO771BR1-GFPLuc+ cells via IC. BMs were allowed to develop for 5 days and mice were randomized in 3 groups: **(i)** E2 (n=24), (ii) EWD (n=24), and (iii) EWD+ Let (n=23). 8 days post-injection each group was divided into two subgroups receiving or not a single WBRT 10 Gy dose and euthanized 14 days later. **D** *Ex-vivo* brain total flux (photon/second) per mouse in irradiated (Left) and non-irradiated cohorts (Right) of C, respectively. Data analysis was Kruskal-Wallis followed by uncorrected Dunn's test or with one-way ANOVA followed by Fisher's LSD test. C. Created in https://BioRender.com. LET= Letrozole.

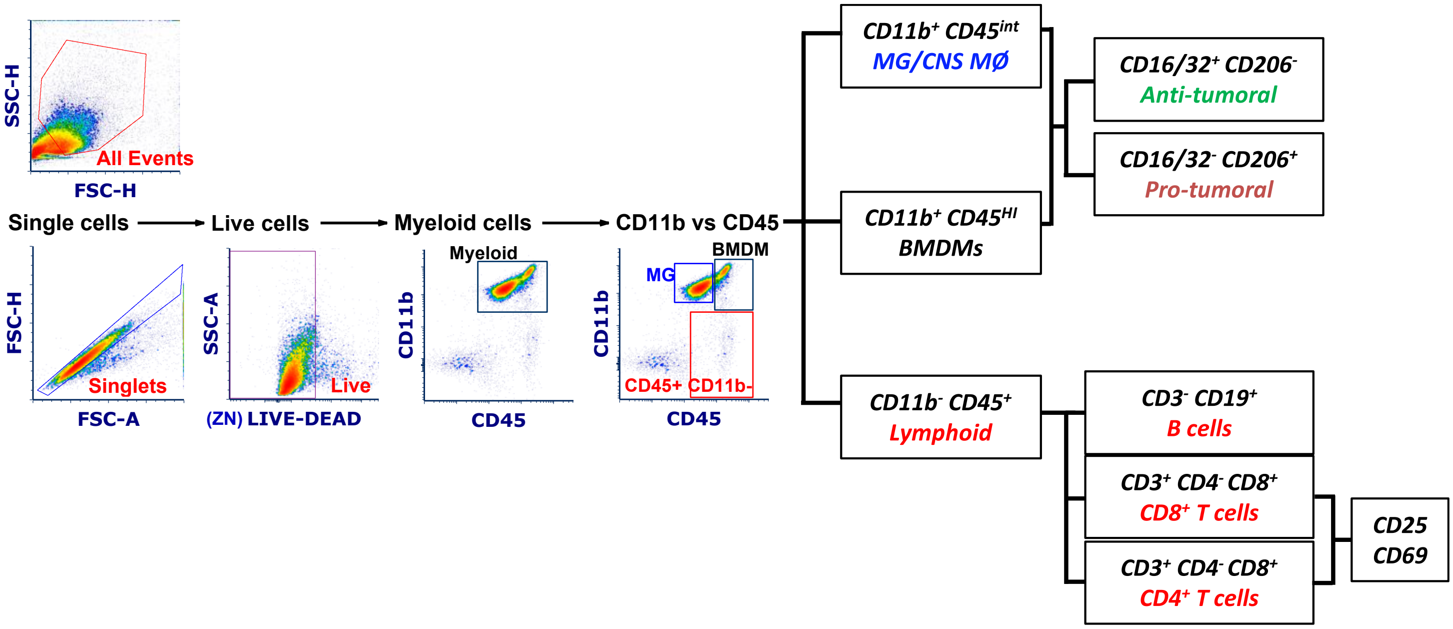


**Supplementary Figure 9.** Manual gating strategy used to define different cell populations in the brain by multiparametric flow cytometry.
