## Supplementary methods for "Estradiol Reprograms Microglia to Create an Immune-Suppressed Niche Permissive to Breast Cancer Brain Metastasis"

**Brain immune cells isolation.** Immune cells from adult mice brains were isolated following the protocol originally outlined by Manglani *et al*[29] with minor adjustments. Briefly, deeply anesthetized mice were transcardially perfused with cold PBS, brains were dissected in ice-cold RPMI phenol red-free media, finely minced with a scalpel and then digested in 1 mg/mL collagenase D (Roche, 11088858001) and 50 µg/mL DNAse (Roche, Cat No. 10104159001) in RPMI phenol red-free media, for 20 min, 37˚C. Digested brains were gently strained through a 100-µm nylon mesh, washed with cold RPMI and centrifuged at 256g (5 minutes at 4°C). Cell pellets were resuspended in 4 mL of cold 90% Percoll and carefully overlayered with 3 mL of 60% Percoll, 4 mL of 40% Percoll, and 3 mL of 1x HBSS. Gradient was then centrifuged at 514 g for 20 minutes at 4°C, with the brake disengaged. After centrifugation, the top 4 mL of the gradient, which includes the myelin layer, was discarded, and the remaining portion was collected, excluding the bottom 4 mL. Cells were rinsed with 25 mL of HBSS and centrifuged at 348 g for 10 minutes at 4°C. The cell pellet was resuspended in 3 mL of PBS, transferred to a 5-ml round-bottom polystyrene tube with a cell strainer snap cap, centrifuged at 400g for 5 minutes and resuspended in 60 uL prior to staining.

**Multiparametric flow cytometry staining:** $1x10^6$ brain or spleen cells were resuspended in 60 µl of PBS and stained with viability stain Zombie NIR at 4°C for 15 minutes. A master mix containing an antibody cocktail was prepared for all samples and diluted in FA3 staining buffer (DPBS calcium and magnesium-free, 10 nM HEPES, 2 mM EDTA, 1% FBS, and 0.1% Sodium Azide). 30 µL of the antibody mix was added to each sample and incubated for 30 minutes in the dark.
