## Supplementary Tables for "Estradiol Reprograms Microglia to Create an Immune-Suppressed Niche Permissive to Breast Cancer Brain Metastasis"

**Table 1:** Antibodies used

| Antibodies/ Fluorochrome | Vendor | Catalog | Clone | RRID | Application |
|---|---|---|---|---|---|
| CD4/BV750 | Biolegend | 100467 | GK1.5 | AB_2734150 | Multiparametric flow cytometry (MFC) |
| CD4/BV510 | Biolegend | 100559 | RM4-5 | AB_2562608 | MFC |
| CD4/AF647 | Biolegend | 100530 | RM4-5 | AB_389325 | MFC |
| CD45/ BV510 | Biolegend | 103138 | 30-F11 | AB_2563061 | MFC |
| CD206/ BV605 | Biolegend | 141721 | C068C2 | AB_2562340 | MFC |
| CD206/BV711 | Biolegend | 141727 | C068C2 | AB_2565822 | MFC |
| CD11b/ BV711 | Biolegend | 101242 | M1/70 | AB_2563310 | MFC |
| CD11b/ PE/Fire 810 | Biolegend | 101285 | M1/70 | AB_2904271 | MFC |
| CD11b/ PE | Biolegend | 101207 | M1/70 | AB_312790 | MFC |
| CD69/ BV785 | Biolegend | 104543 | H1.2F3 | AB_2629640 | MFC |
| CD69/ BV650 | Biolegend | 104541 | H1.2F3 | AB_2616934 | MFC |
| CD8a/ FITC | Biolegend | 100804 | 5H10-1 | AB_312765 | MFC |
| CD8a/ Pacific Blue | Biolegend | 100725 | 53-6.7 | AB_493425 | MFC |
| CD16/32/ PE/CY5 | Biolegend | 156618 | S17011E | AB_2894425 | MFC |
| Ly-6G/ AF647 | Biolegend | 127610 | 1A8 | AB_1134159 | MFC |
| CD19/ AF700 | Biolegend | 115528 | 6D5 | AB_493735 | MFC |
| Ly-6C/ APC/Fire 750 | Biolegend | 128046 | HK1.4 | AB_2616730 | MFC |
| CD3/ APC/Fire 810 | Biolegend | 100268 | 17A2 | AB_2876392 | MFC |
| Ly-6C/PE | Biolegend | 128008 | HK1.4 | AB_1186133 | MFC |
| CD25/PE/Dazzle 594 | Biolegend | 101920 | 3C7 | AB_2721702 | MFC |
| CD16/32/PE/CY7 | Biolegend | 101318 | 93 | AB_2104156 | MFC |
| CD19/AF700 | Biolegend | 115528 | 6D5 | AB_493735 | MFC |
| INFγ/ APC | Biolegend | 505810 | XMG1.2 | AB_315404 | MFC |
| GZMB/ BV650 | Biolegend | 396436 | QA18A28 | AB_3662293 | MFC |

**Table 2:** Antibody panels for multiparametric flow cytometry

| Panel 1 Figure 1 | Panel 2 Figure 2, 4 | Panel 3 Figure 5B-G | Panel 3 Figure 5I-N |
|---|---|---|---|
| CD45: BV510 | CD206: BV711 | CD8a: FITC | CD8a: Pacific Blue |
| CD206: BV605 | CD45: BV510 | CD4: BV510 | CD4: AF647 |
| CD11b: BV711 | CD11b: PE/Fire 810 | CD11b: PE/Fire 810 | CD11b: PE |
| CD4: BV750 | CD4: BV750 | CD69: BV785 | |
| CD69: BV785 | CD69: BV650 | CD25: PE/Dazzel 594 | |
| CD8a: FITC | CD8a: FITC | INFγ: APC | |
| Ly-6C: PE | Ly-6C: PE | GZMB: BV650 | |
| CD25: PE/Dazzle 594 | CD25: PE/Dazzle 594 | | |
| CD16/32: PE/Cy7 | CD16/32: PE/Cy7 | | |
| CD19: AF700 | CD19: AF700 | | |
| Ly6-G: APC/Fire750 | Ly6-G: APC/Fire750 | | |
| CD3: APC/Fire810 | CD3: APC/Fire810 | | |

The cell-specific markers include: CD45 for all immune cells, CD45 and CD11b for microglia/BMDM, CD3, CD4 and CD8 for CD4 T and CD8 T cells, CD19 for B cells.
